# HIV-1 resistance to islatravir/tenofovir combination therapy in wild-type or NRTI resistant strains of diverse HIV-1 subtypes

**DOI:** 10.1101/2023.06.18.545484

**Authors:** Maria E. Cilento, Xin Wen, Aaron B. Reeve, Obiaara Ukah, Alexa Snyder, Ciro M. Carrillo, Cole P. Smith, Kristin Edwards, Claudia C. Wahoski, Deborah R. Kitzler, Eiichi N. Kodama, Hiroaki Mitsuya, Michael A. Parniak, Philip R. Tedbury, Stefan G. Sarafianos

## Abstract

Tenofovir disoproxil fumarate (TDF) and islatravir (ISL, 4’-ethynyl-2-fluoro-2’-deoxyadensine, or MK-8591) are highly potent nucleoside reverse transcriptase inhibitors. Resistance to TDF and ISL is conferred by K65R and M184V, respectively. Furthermore, K65R and M184V increase sensitivity to ISL and TDF, respectively. Therefore, these two nucleoside analogs have opposing resistance profiles and could present a high genetic barrier to resistance. To explore resistance to TDF and ISL in combination, we performed passaging experiments with HIV-1 WT, K65R, or M184V in the presence of ISL and TDF. We identified K65R, M184V, and S68G/N mutations. The mutant most resistant to ISL was S68N/M184V, yet it remained susceptible to TDF. To further confirm our cellular findings, we implemented an endogenous reverse transcriptase assay to verify *in vitro* potency. To better understand the impact of these resistance mutations in the context of global infection, we determined potency of ISL and TDF against HIV subtypes A, B, C, D, and circulating recombinant forms (CRF) 01_AE and 02_AG with and without resistance mutations. In all isolates studied, we found K65R imparted hypersensitivity to ISL whereas M184V conferred resistance. We demonstrated that the S68G polymorphism can enhance fitness of drug-resistant mutants in some genetic backgrounds. Collectively, the data suggest that the opposing resistance profiles of ISL and TDF suggest that a combination of the two drugs could be a promising drug regimen for the treatment of patients infected with any HIV-1 subtype, including those who have failed 3TC/FTC-based therapies.

## Introduction

HIV is a global public health issue that continues to be a threat to public health. In 2020 alone, 680,000 people died from HIV-related causes and 1.7 million were newly infected. HIV-1 can be divided into phylogentic groups (M, N, O, P) resulting from discrete zoonotic transfers (Hemelaar et al., 2006). Of note, group M accounts for the majority of HIV infections. Group M has high genetic diversity and is subdivided into various subtypes: A, B, C, D, F, G, H, K, and L, with new subtypes still being discovered (Taylor & Hammer, 2008). The amino acid sequence variation among the subtypes is typically 20 to 25%, while the intra-subtype variation is between 15 to 20% (Hemelaar et al., 2006; Taylor & Hammer, 2008). In addition, recombinant forms can be generated within a patient infected by two subtypes, resulting into what are known as circulating recombinant forms (CRFs) (Taylor & Hammer, 2008). CF01_AE (originally identified as subtype E) was first identified in Thailand in the late 1980s and dominates southeast Asia (Carr et al., 1996; Gao et al., 1996; McCutchan et al., 1992). CRF02_AG is another common circulating recombinant form found in Western and West Central Africa (Taylor & Hammer, 2008). Of all the subtypes noted, the most studied and well characterized is subtype B (HIV-B), which is primarily found primarily in the US, Europe, and Japan, but accounts for only 11% of global infections (Fettig et al., 2014). HIV-nonB subtypes (such as C, CRF_AE, and CRF_AG) are predominant in Africa and account for the majority of global HIV infections. To implement anti-HIV therapies on a global scale, it is necessary to consider their potency and resistance profiles in the most prevalent HIV subtypes.

HIV-related deaths have decreased over the last 20 years due to highly active retroviral therapy (HAART). HAART typically consists of two nucleoside (or nucleotide) reverse transcriptase inhibitors (NRTIs) and one non-nucleoside reverse transcriptase inhibitor or an integrase inhibitor. The current FDA-approved NRTIs are abacavir, emtricitabine (FTC), lamivudine (3TC), tenofovir (TFV), and zidovudine. The most commonly used NRTIs in the United States are FTC and tenofovir (Schinazi et al., 2022). While current therapies used for the treatment of HIV are efficient in suppressing viral loads, interruptions in patient adherence can lead to the emergence of drug resistant strains, transmission of these strains is a major public health challenge. Hence, there is a continuing need to design new drug combinations. Raising the genetic barrier to resistance can be achieved through various means, such as combining drugs that have different resistance profiles, e.g., because they exploit distinct targets (Lu et al., 2018), or drugs that hit the same target but have opposing resistance profiles, where resistance to one agent confers enhanced sensitivity to the second (Michailidis et al., 2014).

The major clinical resistance mutations to current frontline NRTIs are K65R (TFV resistance mutation) and M184V (FTC and lamivudine (3TC) resistance mutation). In addition, there are several reported subtype-specific differences in treatment response among HIV infected individuals and HAART treated patients. For example, subtype-C patients are more likely to fail on current standard-of-care tenofovir-based regimens, due to higher prevalence in this subtype of the K65R tenofovir resistance mutation in RT (Theys et al., 2013).

Preclinical development of ISL has demonstrated high potency and favorable toxicity profiles in cell culture, mice, and rhesus macaques (Kawamoto et al., 2008; Murphey-Corb et al., 2012; Ohrui et al., 2007; Stoddart et al., 2015). Early clinical studies demonstrated that ISL was well tolerated in humans (Molina et al., 2022). However, in subsequent clinical studies patients on ISL experienced a decline in CD4 T cells, and as a result the US Food and Drug Administration (FDA) placed a hold on ISL clinical trials(Merck & Co., 2021). Recently, new clinical trials on ISL-based therapies were announced, as it was shown that a lower dose of ISL maintains the benefits without any effects on the CD4 T cell population (Merck & Co., 2022).

ISL has three structural features that contribute to its efficacy and long-acting potential. Unlike currently approved HIV NRTIs, ISL retains a 3’-OH, which enhances efficient activation by the cellular deoxycytidine kinase to create ISL-monophosphate (ISL-MP), the first and rate-limiting activation step of almost all NRTIs (Kawamoto et al., 2008). The 2-fluoro (2-F) imparts resistance to adenosine deaminase, a key metabolizing enzyme of adenosine and adenosine-analogues (Kirby et al., 2013). The 4’-ethynyl (4’-E) contributes to the exceptional binding affinity of ISL at the reverse transcriptase (RT) polymerase active site through interactions with residues in a conserved hydrophobic pocket: A114, Y115, F160, and M184, and the aliphatic part of D185 (Salie et al., 2016). ISL inhibits HIV-1 RT by at least two mechanisms: a) immediate chain termination (ICT), where ISL-MP is incorporated then blocks DNA synthesis by inhibiting translocation; or b) delayed chain termination (DCT) where ISL-MP is incorporated and followed by a single additional nucleotide prior to chain termination (Michailidis et al., 2014; Michailidis et al., 2009). Also, previous studies with ISL have shown that K65R hypersensitizes RT to ISL (4-fold more susceptible than wildtype RT), whereas M184V confers mild resistance (8-fold relative to wildtype RT) (Michailidis et al., 2009).

Here, we explore how pre-existing resistance mutations affect the emergence of resistance to tenofovir and ISL combination regimens. We began passaging experiments with WT, K65R (resistance mutation in RT to tenofovir), and M184V (resistance mutation to 3TC/FTC and ISL). To confirm the resistance findings, both cell-based and endogenous reverse transcriptase assays were used to determine sensitivity to ISL and TFV. We also explored clinical resistance of K65R, M184V, and K65R/M184V in the context of diverse primary isolates: B, C, CRF_AE, and CRF_AG. In all isolates studied, we found K65R imparted hypersensitivity to ISL. M184V confers resistance to ISL that ranges from 3- to 10-fold relative to wildtype. The K65R/M184V double mutant typically maintained or decreased resistance to ISL compared to the M184V mutant alone. We further found that S68G, when combined with K65R/M184V, in some genetic contexts, conferred greater resistance than M184V alone. S68G can emerge during TVF-treatment ^34^. Collectively, the data suggest that ISL and TDF is a promising drug regimen for the treatment of HIV-1 due to their opposing resistance profiles, high barrier to resistance, and capability of suppressing a wide range of HIV-1 subtypes.

## Methods

### Reagents

Stock solutions of ISL (Life Chemicals, Canada) and TDF (provided by AIDS Reagent Program) were prepared in distilled, deionized H_2_O. MT-2 cells (Charneau et al., 1994; Haertle et al., 1988) were cultured in RPMI 1640 (Mediatech, Inc., Manassas, VA), supplemented with 10% fetal bovine serum (FBS; HyClone, Logan, UT) and 2 mM L-glutamine, 100 U/mL penicillin/streptomycin (Thermo Fisher). (Mediatech, Inc.). HEK-293/17 (Pear et al., 1993) and TZM-GFP cells from Massimo Pizzato (Trento University) cells were cultured in DMEM (Corning) supplemented with 10% Serum Plus II (Sigma-Aldrich, St. Louis, Missouri), 2 mM L-glutamine, and 100 U/mL penicillin/streptomycin.

### Generation of virus stocks and molecular clones

K65R and M184V mutations in RT were generated by site-directed mutagenesis on an HXB2 or LAI HIV-1 backbone using the QuikChange XL site-directed mutagenesis kit (Agilent Technologies, Inc., Santa Clara, CA) according to the manufacturer’s protocols. 293-T cells were transfected with 10 μg of plasmid DNA using the PrimeFectimine mammalian transfection reagent (PrimGen, Oak Park, IL). After 72 h incubation, media were harvested, filtered, and used to infect MT-2 cells and infectious virus harvested at ≥50% syncytium formation. HIV-1 infectious clones, containing the gag/pol coding regions of primary isolates into an NL4-3 backbone, were obtained from Drs. Ujjwal Neogi and Anders Sönnerborg. The Cloning Core of Emory University introduced putative resistance mutations into these primary isolate-derived infectious clones. Virus stocks were made using HEK-293/17 cells that were transfected with 5 μg of viral DNA using X-tremeGENE HP (Roche, Basel, Switzerland) after 48 h incubation, HEK-293/17 cell media were harvested and concentrated overnight with a Lenti-X concentrator (Clontech, Mountain View, CA) according to the manufacturer’s protocol and resuspended in 10-fold less media volume prior to concentration.

### Serial passage for selection of resistant virus

MT-2 cells were seeded at 2.5 x 10^6^ cells per 1 mL in 10 mL of media containing ISL and TDF. Initial ISL concentrations were chosen based on the EC_50_ of wildtype stock virus. The unpassaged WT stock virus used in these passages had an ISL EC_50_ of 3.6 nM. For TDF, the WT stock had an EC_50_ of 24.6 nM. The TDF:ISL EC_50_ value ratio was determined to be 0.6:1, 7:1 and 80:1 for M184V, WT and K65R, respectively. For simplicity, we elected to passage each virus in 1:1, 10:1 or 100:1 TDF:ISL combinations. Every 2-3 days the cells were mixed to ensure even suspension of cells and 9 mL of the media was removed and replaced with fresh media containing the appropriate concentration of ISL and TDF. At ≥ 75% syncytia formation, culture supernatants were harvested, concentrated using Amicon Ultra Ultracel – 100,000 MWCO centrifugal filters (Millipore, Carrigtwohill, Co. Cork, Ireland) and syringe-filtered through 0.22 µM filters (Millipore).

### Sequencing of passaged virus

Viral RNA was purified from supernatants using the QIAamp Viral RNA Mini kit (Qiagen, Valencia, CA), the concentration determined using Spectronic BioMate*3 UV spectrophotometer (Thermo Scientific) and 500 ng RNA was used as the template for cDNA synthesis. First-strand PCR was performed using random hexamer primers and the SuperScript III First-Strand Synthesis System for RT-PCR (Invitrogen, Carlsbad, CA). The resulting cDNA was PCR amplified using HIV-1 LAI-specific primers ABR-RT-OF (1763 5’-GGAGCCGATAGACAAGGAACTG-3’) and ABR-RT-OR2 (3863 5’-GGCTACTATTTCTTTTGCTACTACAGG-3’). These primers are located in the 3’ end of gag and the 5’ end of integrase, respectively, and generate a 2127 bp product spanning the full length of the reverse transcriptase gene. PCR was performed using the Expand High Fidelity PCR System dNTPack (Roche Diagnostics GmbH, Mannheim, Germany).

Approximately 100 ng of full-length PCR product was ligated into the pGEM-T Vector System (Promega, Madison, WI), at a 3:1 molar ratio of insert: vector and incubated overnight at 4°C.Ligations were transformed into MAX Efficiency DH5α competent cells (Invitrogen, Carlsbad, CA). Blue-white screening was used to select clones with successful ligations, and plasmids containing the full-length reverse transcriptase gene were isolated with the QIAprep Spin Miniprep kit (Qiagen). Primers ABR-RT-OF and ABR-RT-OR2 were used to sequence the 5’ and 3’ ends of reverse transcriptase, while an internal portion of the gene was sequenced with primer ABR-RT-IF (2211 5’-CAGAGATGGAAAAGGAAGGG-3’). Clones were aligned with the appropriate P0 stock virus consensus in Clustal X-2, and the proportion of sequences with the novel substitutions was determined.

### Dose-response assays to determine sensitivity to antivirals

After each virus was generated as described above, TZM-GFP cells were plated at 10,000 cells/well in a 96-well plate and with serially diluted ISL or TDF at 0.000381 nM to 100 nM (ISL) or 0.038 nM to 10 μM (TDF). The cells, media, and drug were incubated for 24 h and infected with virus and a 1 μg/mL final concentration of DEAE-dextran, followed by incubation for 48 h. The green fluorescent protein (GFP)-positive cells were then counted using Cytation 5 with Gen5 v3.03 software (BioTek, Winooski, VT). Dose-response curves were plotted and EC_50_ values determined using Prism 9 (GraphPad) software.

### Determination of specific infectivity

TZM-GFP cells were plated at 10,000 cells/well in a 96-well plate and incubated for 24 h. Then, cells were infected with the serially diluted virus and a 1 μg/mL final concentration of DEAE-dextran, followed by incubation for 48 h. The GFP-positive cells were then counted as described above. The p24 content of each virus stock was also determined using an enzyme-linked immunosorbent assay (ELISA). Specific infectivity was calculated as infected cells divided by the amount of p24 content.

### Endogenous reverse transcriptase assay

RT protocols were adapted from (Christensen et al., 2020; Jennings et al., 2020) reactions were performed in 50 μL containing 10 mM Tris-HCl, pH 7.8, 75 mM NaCl, 2 mM MgCl_2_, 2.5 μg/mL melittin, 80 µM IP6, 0.004 mM dCTP, dTTP, and dGTP, 0.02 μM dATP, 100 ng capsid, and RT inhibitors, including ISL-TP and Tenofovir-DP. After 30 minutes incubation at 37°C, 0.036 mM of dCTP, dGTP, dTTP, and 0.0398 mM dATP were added to the reaction and incubated at 37°C for a further 16 h. The products were quantified by qPCR with SYBR green detection using these primers: 5’-GAGCTCTCTGGCTAACTAG-3’ and 5′-TGACTAAAAGGGTCTGAGGGATCT-3′ adapted from ^41^.

### Statistics

The statistical significance was determined using Prism 9. A one-way analysis of variance (ANOVA) was performed, as well as a Dunnett’s or Tukey’s multiple-comparison test (see figure legends).

## Results

### Resistance mutations found during serial passages of WT, K65R, M184V viruses in the presence of ISL and TDF

Serial passages of HIV-1 in MT-2 cells were performed with WT, K65R, and M184V viruses to identify mutations conferring resistance to ISL and TDF combination treatment. The WT virus survived six passages in 1:1 TDF:ISL (increasing drug concentration by 2-fold for each passage), escaping from final concentrations of 14.4 nM TDF and 14.4 nM ISL, over the course of 34 days. M184I was detected in 71.4% of the passage 6 (P6) sequences and M184V in 28.6%. When the TDF:ISL ratio was increased to 10:1, the WT virus escaped from only five passages over 65 days, with final drug concentrations of 144 nM TDF and 14.4 nM ISL; M184V was present in 100% of the final passage (Table 1). At 100:1 TDF:ISL (close to 1:1 when comparing EC_50_ values), the WT virus required 97 days to escape two passages and could only survive the initial combination of 360 nM TDF and 3.6 nM ISL. The virus isolated had no mutations except M184I at 9.5% (Table 1).

**Table 1.**
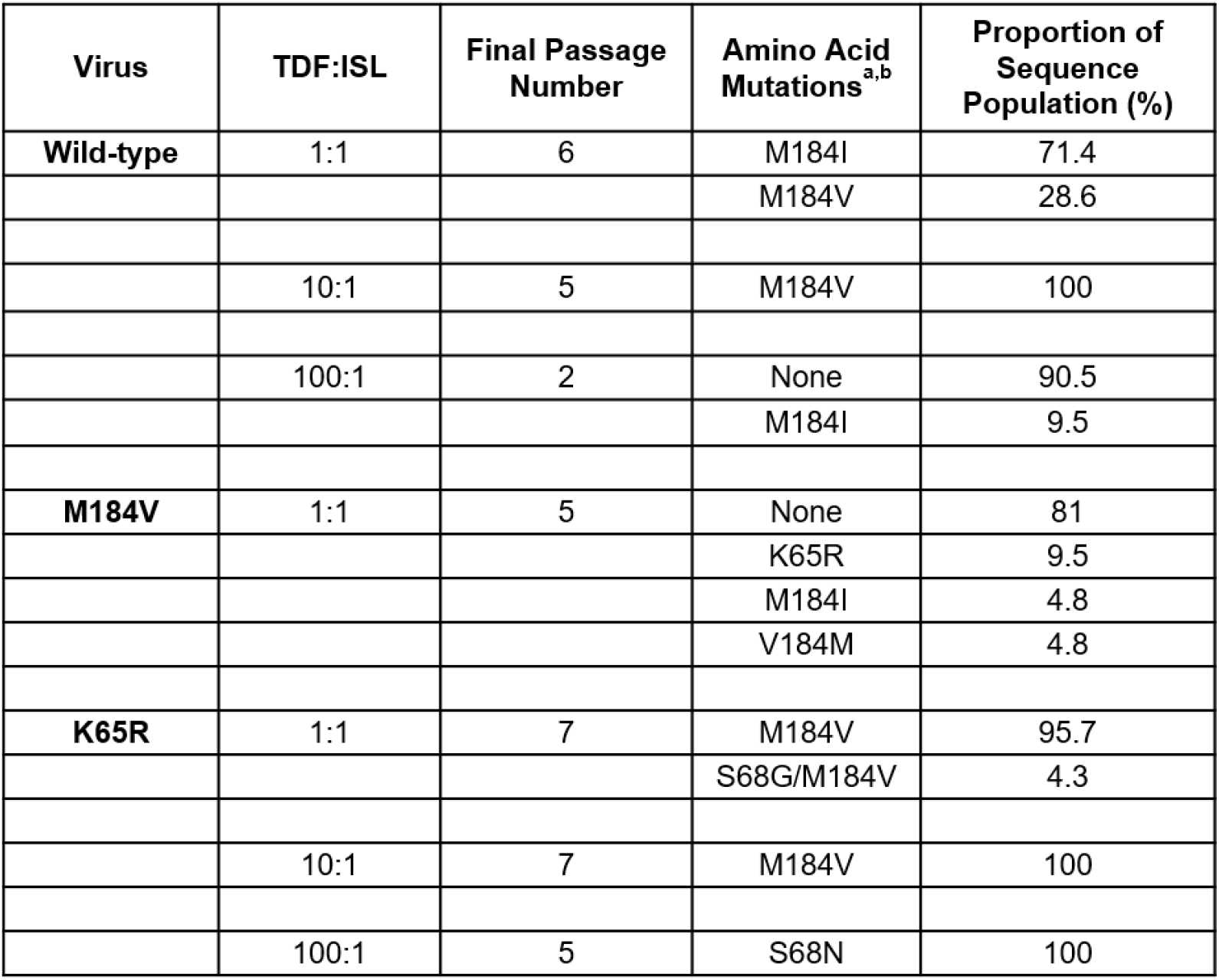
Amino acid mutations identified in wildtype, K65R and M184V viruses following serial passage in TDF:ISL.

The M184V strain progressed through five passages in 1:1 TDF:ISL, spanning 87 days and reaching final concentrations of 130 nM TDF and 130 nM ISL, where K65R/M184V was found at 9.5%, M184I was 4.8% and a reversion to M184 was also found in 4.8%. M184V was unable to escape from even a single passage in 10:1 or 100:1 TDF:ISL.

The K65R virus survived seven passages in 1:1 TDF:ISL over 32 days, escaping from a final combination of 7.2 nM TDF and 7.2 nM ISL. K65R/M184V took over the final population of 95.7% at passage 7. K65R/S68G/M184V was found at 4.3% prevalence at passage 7. In 10:1 TDF:ISL, the K65R strain progressed through seven passages over 36 days and escaped a final combination of 72 nM TDF and 7.2 nM ISL. K65R/M184V appeared in 100% of the population (Table 1). The K65R virus only survived five passages in 100:1 TDF:ISL over 58 days, emerging from 360 nM TDF and 3.6 nM ISL where K65R/S68N was found in 100% of the populations. Regardless of the starting virus (WT, K65R, and M184V), M184V/I was consistently found except when there was 100:1 ratio of TDF:ISL.

### Validation of mutations found in TDF- and ISL-passaging using molecular clones and evaluating the specific infectivity

To evaluate the contributions of the individual mutations, and combinations of mutations, identified during viral passaging experiments, we introduced mutations into the HIV-1 NL4-3 lab strain and determined sensitivity to ISL and TDF (Figure 1A). M184V conferred 7-fold resistance to ISL and was 2-fold more sensitive to TDF than WT, consistent with previous reports ^19,43-46^. K65R was found to be hypersensitive to ISL (4-fold more sensitive than WT) and 3-fold resistant to TDF, which was also consistent with previous findings (Brenner & Coutsinos, 2009; Cilento et al., 2021; Deval et al., 2004; Kawamoto et al., 2008; Maeda et al., 2014; Margot et al., 2002; Michailidis et al., 2013; Miller, 2004). The K65R/M184V double mutant was more sensitive to ISL that M184V alone (3-fold resistance relative to wildtype compared to 7-fold for M184V), and more sensitive to TDF than K65R alone (2-fold resistance relative to wildtype compared to 4-fold for K65R), but the double mutant also lacked the hypersensitivities of the single mutants. In that respect, viruses containing K65R/M184V may be advantageous for the viruses that are passaged under selection of both ISL and TDF. The additional mutants, S68G/M184V and S68N/M184V, had 6-fold and 12-fold resistance to ISL, respectively, while both were found to be susceptible toTDF (Figure 1A). No major resistance to ISL (>12-fold resistance) or TDF (>4-fold resistance) was observed.

**Figure 1.**
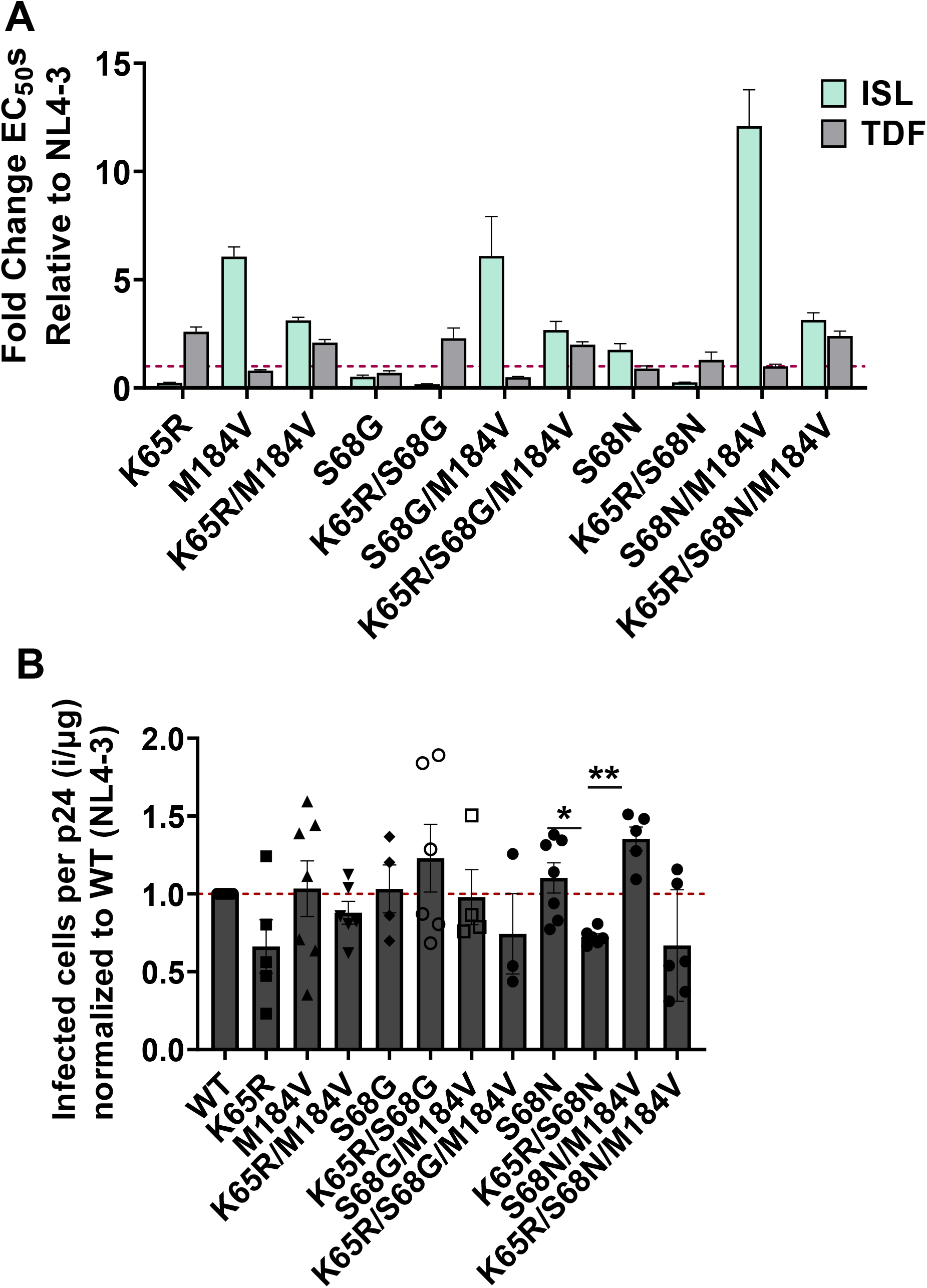
Resistance and specific infectivity of mutants identified during ISL and TDF passaging. (A) Potency of ISL and TDF mutants. TZM-GFP cells were pretreated with serial dilutions of ISL or TDF then infected with HIV-1. 48 h post infection, infected cells were counted. Dose-response curves were produced for each mutant and the EC50s calculated. EC_50_s were expressed as fold change from WT. Dashed line denotes NL4-3. Statistical significance was determined using one-way ANOVA with Tukey’s posttest (***, P < 0.001). The results represent the means and SD for three independent experiments performed in duplicate. (B) Specific Infectivity of mutants. Single-round infection of TZM-GFP cells was performed withHIV-1 mutants, following normalization of input p24 by ELISA. The ratio of infected cells per normalized input (p24) was then calculated and expressed relative to NL4-3. Dashed line denotes NL4-3 Statistical significance was determined using a one-way ANOVA with Dunnett’s multiple-comparison test (*, *P <* 0.05; **, *P <* 0.01). The results represent the means and SD for three to six independent experiments in triplicate.

To determine whether the mutations identified had effects on viral fitness, we used a single round replication assay to determine the specific infectivity of viral stocks. Specific infectivity was determined by amount of infection per amount of p24 or capsid. The only significant changes in specific infectivity compared to WT were S68N/M184V and K65R/S68N, which had a minor increase and decrease, respectively (Figure 1B).

To further verify the effects of ISL and TDF resistant mutations, removed from the context of infection in the context of endogenous RT, we implemented an endogenous RT assay. We used isolated virions supplemented with dNTPs, IP6, and phosphorylated ISL (ISL-triphosphate, ISL-TP) and tenofovir (TFV-diphosphate, TFV-DP), and RT products using qPCR. The resistance phenotypes broadly recapitulated those observed in cell-based assays (Figure 1A and 2); K65R and M184V conferred resistance to TFV-DP and ISL-TP, respectively. There was one notable deviation in the data, we were unable to determine IC50 values for mutants containing both K65R and M184V. Manual inspection of the raw data suggested elevated resistance to both antivirals, but data were too variable to permit a reliable dose-response curve to be plotted.

**Figure 2.**
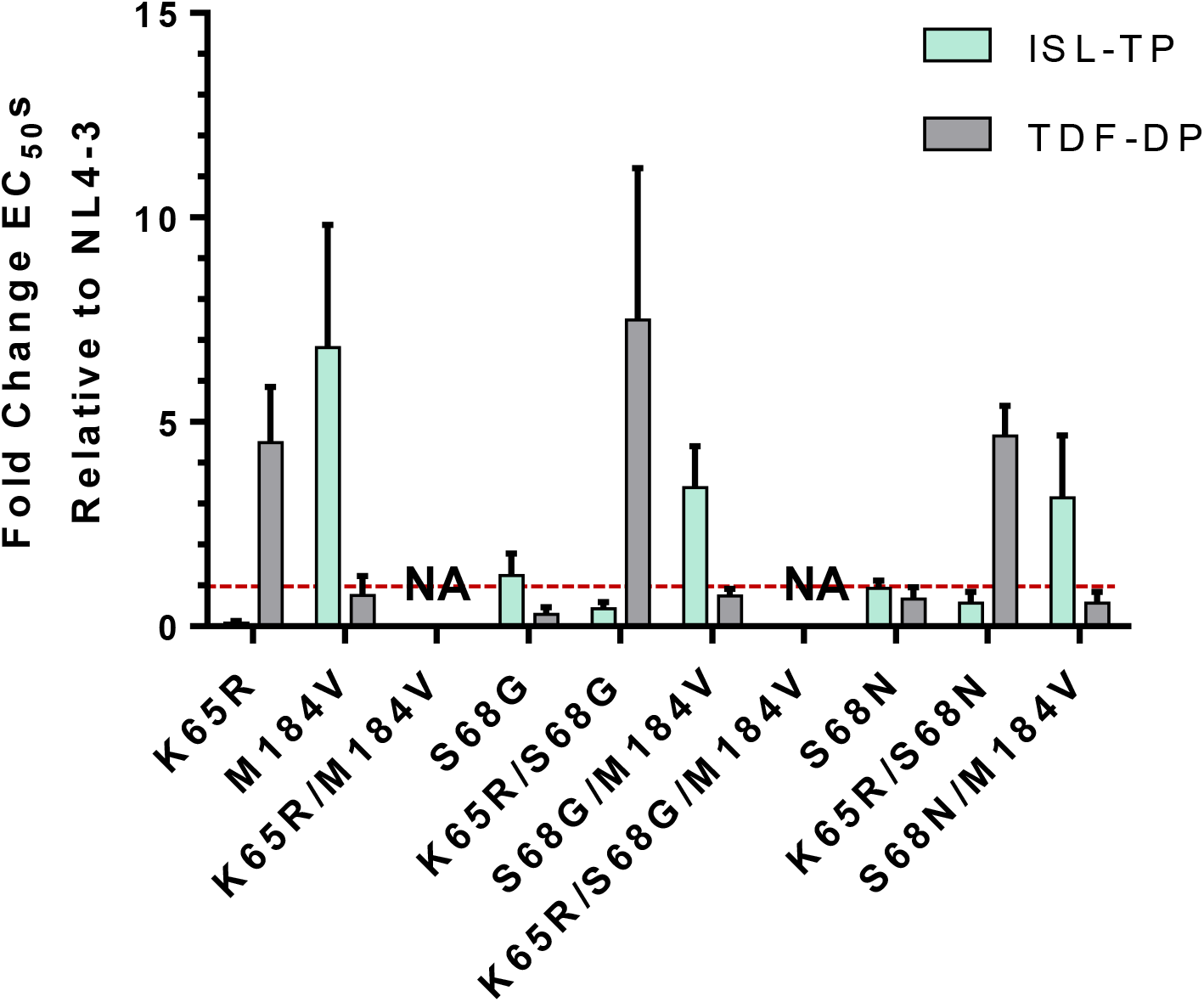
Potency of ISL and TFV against resistant mutants *in vitro*. Viral particles were produced by transfection of mutant infectious clones into 293T cells. Virions were harvested and samples normalized by ELISA. RT activity was measured by endogenous RT assay in the presence or absence of antivirals. Dose-response curves were produced for each mutant and the IC_50_s calculated. Mutants were normalized to WT infection to produce the fold change. Dashed line denotes NL4-3. The results represent the means and SD for two or three independent experiments performed in duplicate. NA, these values could not be calculated from the experimental data.

### Potency of ISL and TDF against diverse HIV subtypes

To determine if TDF and ISL have the potential to be effective on a global scale, we performed dose-response experiments with 8-10 isolates from subtypes A, B, C, D, CRF_AE, and CRF_AG. ISL was found to be broadly effective against subtypes with no more than 1.3-fold change from NL4-3 (Figure 3). Combining all isolates, the mean EC_50_ of ISL was 0.80 nM. TDF was also effective at inhibiting subtypes with a mean EC_50_ of TDF of 87 nM; however, subtypes A, B, C, and D had 1.3-, 2.2-, 2.2-, 1.6-fold change from NL4-3, respectively (Figure 3).

**Figure 3:**
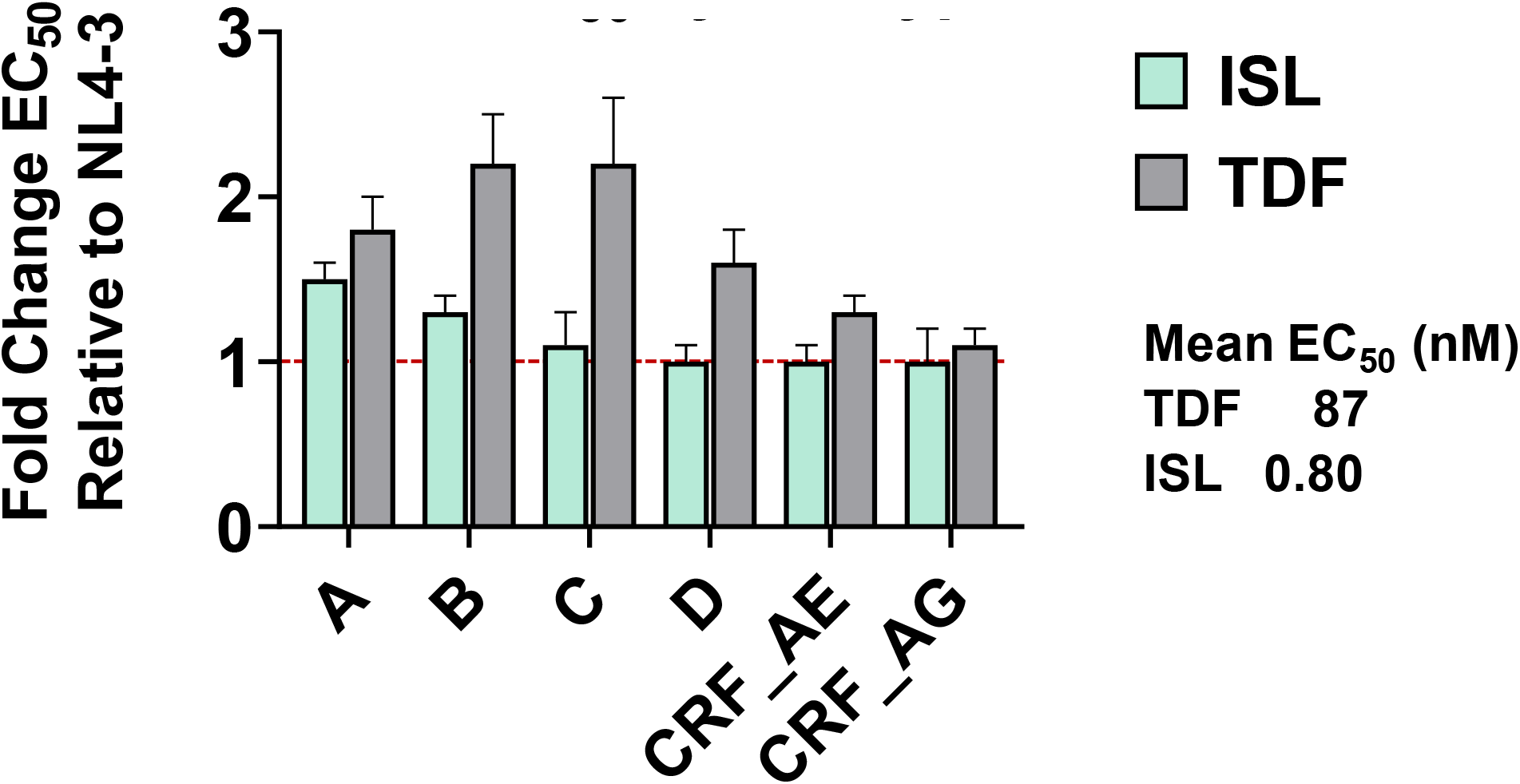
Breadth of activity of ISL and TDF against diverse primary isolates. TZM-GFP cells were pretreated with serial dilutions of ISL or TDF then infected with HIV-1. 48 h post infection, infected cells were counted. Dose-response curves were produced for each mutant and the EC_50_s calculated. EC_50_s were expressed as fold change from WT. Dashed line denotes NL4-3. EC_50_s were determined for 8-10 isolates from each subtype.

### Impact of mutations K65R, M184V, and K65R/M184V on susceptibility to ISL and TDF and viral fitness

To better understand utility for global implementation of TDF/ISL, we examined resistance mutations in the context of diverse primary HIV isolates, *gag/pol* was cloned from isolates B, C, CRF_AE, and CRF_AG into NL4-3 and the K65R, M184V, or K65R/M184V changes introduced in these contexts. Regardless of the isolate studied, K65R conferred hypersensitivity to ISL and 2- to 4-fold more resistant to TDF than WT (Figure 4A). M184V conferred resistance to ISL, and the degree of resistance was found to vary substantially between isolates: CRF_AE, B, C, and CRF_AG exhibited 2.5-fold, 3.5-fold, 9.5-fold, and 8-fold resistance, respectively (Figure 4B). Meanwhile, M184V was found to be as susceptible to TDF as WT regardless of the isolate. Finally, K65R/M184V conferred mild resistance to both ISL and TDF without enhanced sensitivity to either compound (Figure 4B).

**Figure 4.**
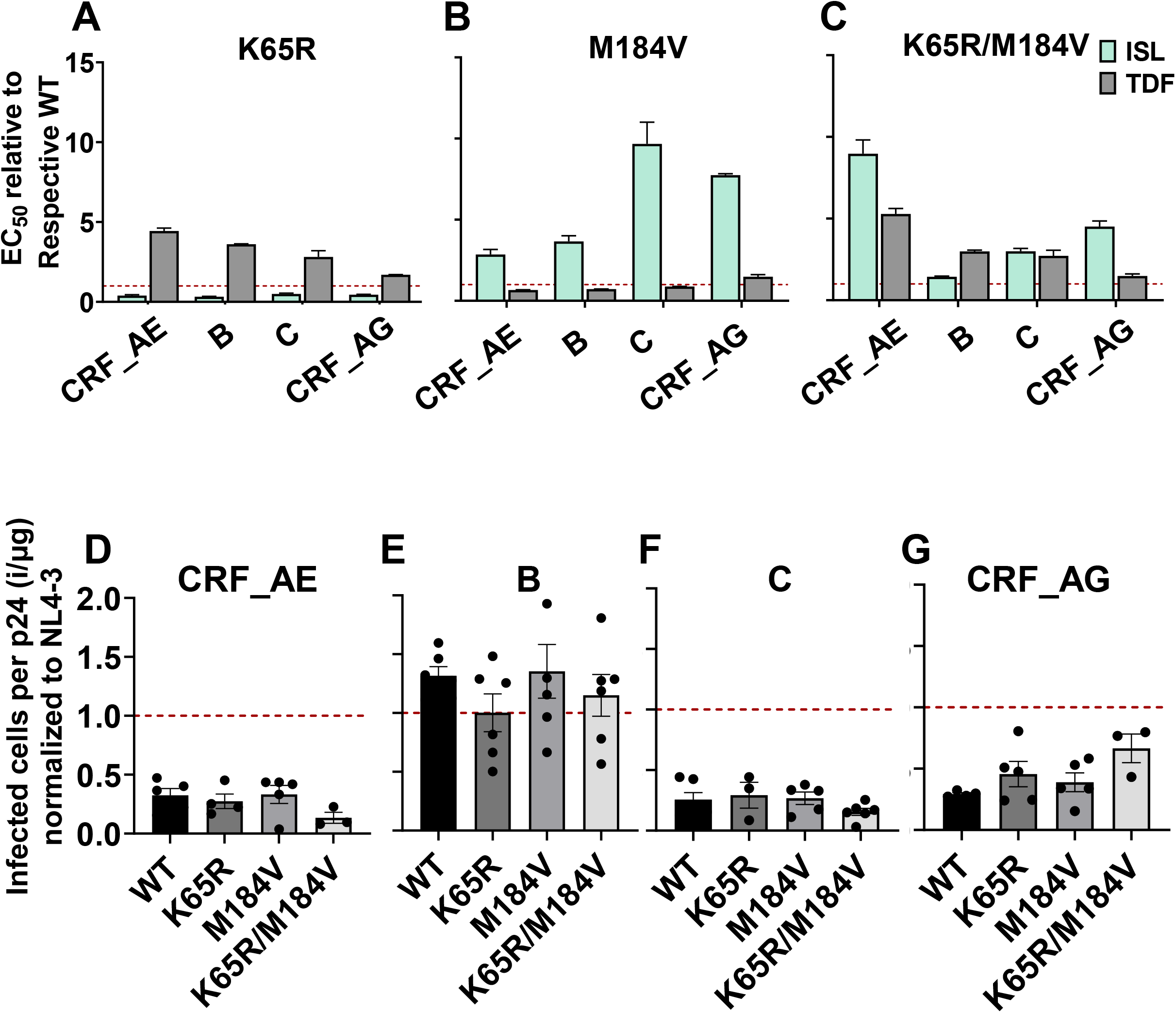
Efficacy of ISL and TDF against clinical isolate RT mutants. (A-C) TZM-GFP cells were pretreated with ISL or TDF and infected with the indicated viruses (A) K65R, (B) M184V, (C) K65R/M184V. 48 h post infection infected cells were counted, dose-response curves were plotted and the EC_50_s expressed relative to their respective WT. Dashed line denotes NL4-3 WT. Statistical significance was determined using one-way ANOVA with Tukey’s posttest (***, P < 0.001). The results represent the means and SD for three independent experiments performed in duplicate. (D-G) Specific Infectivity of clinical isolates containing K65R, M184V, or K65R/M184V mutations. TZM-GFP cells were infected with p24 ELISA-normalized mutant viruses. The ratio of infected cells per p24 was calculated and expressed relative to NL4-3. Dashed line denotes NL4-3. Statistical significance was determined using a one-way ANOVA with Dunnett’s multiple-comparison test. No significant differences relative to the respective WTs were observed.

Unlike the other subtypes, isolate CRF_AE K65R/M184V had greater resistance (9-fold) to ISL than M184V (3-fold). This was unexpected as K65R/M184V double mutants displayed an intermediate phenotype in all other isolates examined, presumably as a consequence of the enhanced resistance conferred by M184V and the enhanced sensitivity of K65R (Figure 4C). B, C, and CRF_AG containing K65R/M184V had 1.4-, 3-, and 4.5-fold resistance to ISL, respectively (Figure 4C). In addition, CRF_AE K65R/M184V had 5.2-fold resistance to TDF, greater than that observed for B, C, and CRF_AG K65R/M184V, which had 3, 2.7-, and 1.5-fold resistance to TDF, respectively. To understand whether these mutations impact the specific infectivity of the isolates, we performed single-round replication viral fitness studies. In CRF_AE, C, and CRF_AG isolates, all mutants had reduced specific infectivity relative to NL4-3 but the mutants had no significant change in specific infectivity compared to their respective WT (Figure 4D, 4F, and 4G). The subtype B isolate had similar specific infectivity to NL4-3 and the mutants K65R, M184V, and K65R/M184V also had no effect on specific infectivity (Figure 4E).

### Polymorphism of Gly68 in CRF_AE K65R/M184V influences susceptibility to ISL and TFV and viral fitness

Inspection of the CRF_AE sequence revealed Gly68, rather than the Ser68 present in the other isolates and the majority of isolates circulating globally. A S68G/N mutation also emerged during passaging experiments with ISL and TDF combinations (Table 1). To test the contribution of residue 68 to drug resistance and fitness, we mutated the Gly68 in CRF_AE back to Ser68 and performed dose-response studies. In the context of the CRF_AE backbone, Ser68 was more susceptible to ISL than Gly68 (Figure 5A). Similarly, K65R with Ser68 was 3-fold more susceptible to ISL than K65R with Gly68. M184V was 3.5-fold resistant to ISL, relative to wildtype, with Ser68 or Gly68 (Figure 5A). In the context of K65R/M184V, Gly68 increased resistance to ISL from 3.5-fold, relative to wild type, to 9-fold resistance, thus contributing to more resistance.

**Figure 5.**
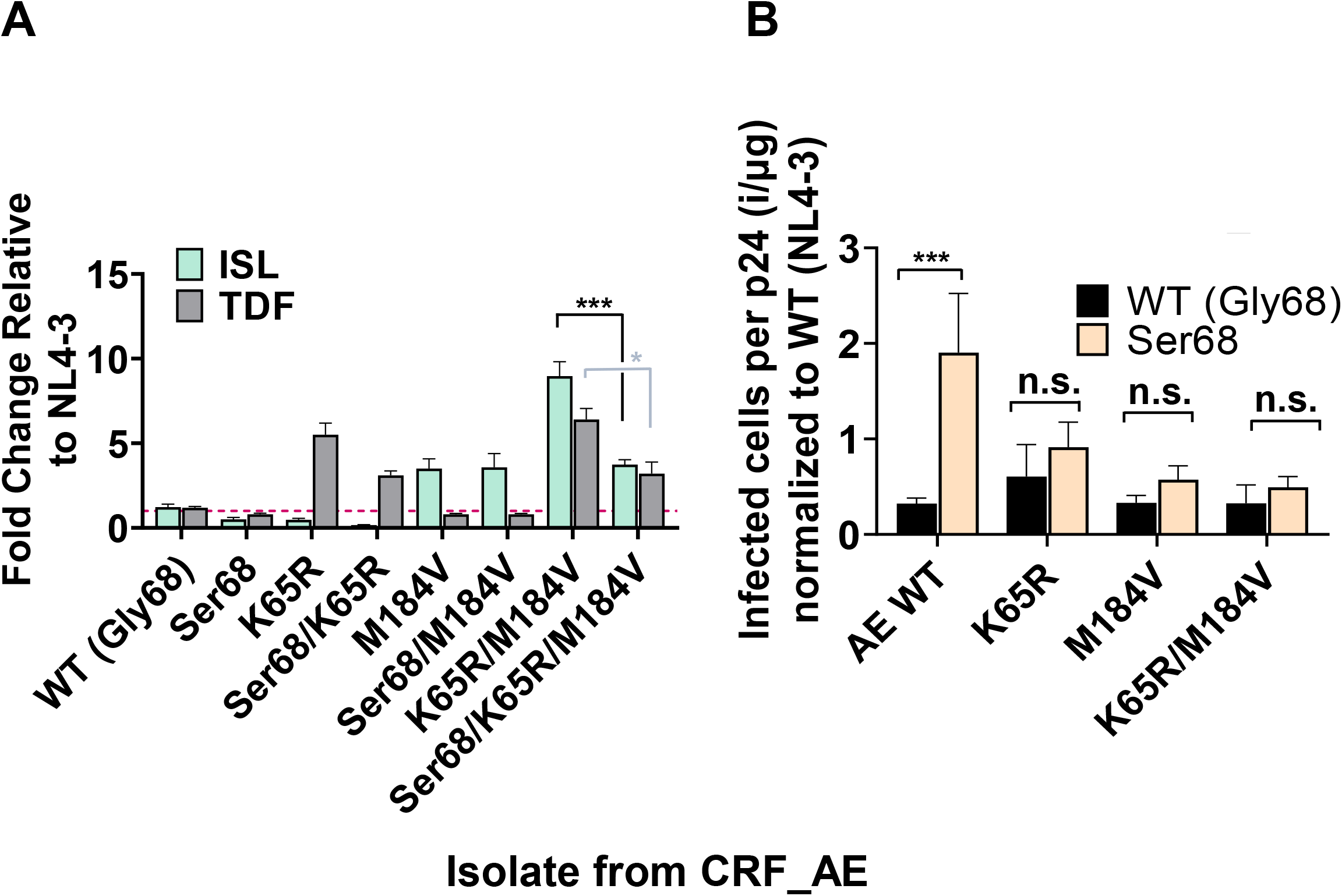
Impact of RT residue 68 mutations on NRTI susceptibility and fitness of CRF_AE. (A) TZM-GFP cells were pretreated with ISL or TDF and after 24 h infected with CRF_AE bearing the indicated mutations. 48 h post infection GFP-positive cells (infected cells) were counted. Dose-response curves were produced and the EC_50_s calculated and expressed as fold-change relative to WT. Dashed line denotes NL4-3 WT. Statistical significance was determined using one-way ANOVA with Tukey’s posttest (***, P < 0.001). The results represent the means and SD for three independent experiments performed in duplicate. (B) Specific Infectivity of CRF_AE mutants indicated was determined by single-round infection assays using TZM-GFP cells infected with ELISA normalized input virus. The ratio of infected cells per p24 was then calculated and normalized to NL4-3. Dashed line denotes NL4-3 Statistical significance was determined using a one-way ANOVA with Dunnett’s multiple-comparison test (***, P < 0.001).

‘To evaluate if Gly68 affects fitness of the CRF_AE isolate, we performed specific infectivity assays. Gly68 reduced specific infectivity in the context of WT CRF_AE, but when combined with other mutations (K65R, M184V, or K65R/M184V) it played no role in specific infectivity (Figure 5B).Some prior studies highlight that S68G is a compensatory mutation associated with K65R (Scherrer et al., 2012; Svarovskaia et al., 2008), while others suggest a polymorphism that does not play a functional role or that it is not linked to K65R at all (Wirden et al., 2005). It was identified here in viral populations with some of the highest resistance to ISL/TDF combination treatment. To better understand the distribution of these populations in the clinic we consulted the HIV Drug Resistance Database maintained by Stanford University (Rhee et al., 2003; Rhee et al., 2006; Shafer, 2006; Shafer et al., 2000). As seen in Table 1, the S68G/N mutations seem to be increased compared to untreated individuals. Taking the Stanford Database, previous studies, and our data S68G/N mutations appear to be slightly enriched in NRTI- and NNRTI-experienced populations (Table 2), consistent with a moderate role in antiviral resistance or viral fitness.

**Table 2.** HIV Drug Resistance Database by Stanford University S68G/N isolate prevalence. Database references: (Rhee et al., 2003; Rhee et al., 2006; Shafer, 2006; Shafer et al., 2000)

| Isolate found | Untreated | Treated with one NRTI | NNRTI treated patients |
| --- | --- | --- | --- |
| S68G | 4% | 4.9% | 8% |
| S68N | 0.24% | 0.44% | 1.4% |

## Discussion

NRTIs have been fundamental in HAART (Lu et al., 2018), but due to their wide use, NRTI-resistance has become increasingly prevalent, albeit the trends vary geographically and can be lacking in the populations of greatest need (Chimukangara et al., 2019; Li et al., 2016; McCluskey et al., 2018; Mega, 2019; Rhee et al., 2020). ISL and TDF hold promise as a combination therapy because of their opposing resistance profiles: K65R confers resistance to TDF (Bazmi et al., 2000; Brenner & Coutsinos, 2009; Miller, 2004; Naeger & Struble, 2006; Zhang et al., 1994) but increased susceptibility to ISL (Kawamoto et al., 2008; Maeda et al., 2014; Michailidis et al., 2013), while M184V confers resistance to ISL (Kawamoto et al., 2008; Maeda et al., 2014) but increased susceptibility to TDF. This relationship between resistance to ISL and sensitivity to TFV is reinforced by a highly (25-fold) ISL-resistant mutant we identified previously, A114S/M184V, which exhibits even more pronounced (40-fold) sensitivity to TDF (Cilento et al., 2021; Diamond et al., 2022). As such, we were interested in identifying and characterizing resistance mutations that might arise with ISL/TDF combination therapy during infection with WT HIV-1 or current NRTI-resistant viruses (K65R or M184V). Importantly, although these two nucleoside analogs are adenosine-based antivirals, they have been shown not to act antagonistically (Hachiya et al., 2013; Michailidis et al., 2014), hence, detailed examination of the interactions between mutations conferring resistance to ISL/TDF combination therapy could be informative for clinical use.

M184V/I confers very high resistance to 3TC/FTC (>100-fold resistance) (Petrella et al., 2004; Sarafianos et al., 1999; Schinazi et al., 1993; Tisdale et al., 1993), but only modest resistance to ISL (∼8-fold resistance) (Kawamoto et al., 2008; Maeda et al., 2014). During viral passage, regardless of the starting virus (WT, K65R, and M184V), M184V/I consistently emerged under conditions where ISL was the dominant antiviral in the combination. In passages that began with K65R present, even low concentrations of TDF were sufficient to maintain K65R in populations, despite the fitness cost that mutation imparts. This is contrary to passaging in the presence of ISL alone, where K65R rapidly reverts to WT (Cilento et al., 2021). In addition, S68G/N was identified in some genetic backgrounds, and was found to enhance resistance and fitness. S68G/N mutations appeared to be associated with K65R, consistent with one previous study (Margot et al., 2006). S68G appears frequently with K65R (Miller, 2004; Roge et al., 2003; Svarovskaia et al., 2008) or Q151M (Garcia-Lerma et al., 2000; Scherrer et al., 2012), a resistance mutation to AZT and other NRTIs (Kavlick et al., 1998; Shirasaka et al., 1995), but not TFV and 3TC/FTC. Another study that sequenced HIV-1 subtype C viruses isolated from 23 patients in Botswana treated with didanosine-based regimens study found that S68G was present with K65R in 3 out of 23 patients (Doualla-Bell et al., 2006). Our results are indicative of a high barrier of resistance to ISL/TDF in combination. It is also important to note that a study with ISL treated rhesus macaques infected with SIV had the M184V mutation present, but was fully suppressed, indicating it was unable to breakthrough ISL therapy (Murphey-Corb et al., 2012).

Previous studies demonstrated that ISL resistance mediated by M184V/I is conferred by steric hinderance of the β-branched amino acids (Val or Ile), which may perturb the hydrophobic pocket into which the 4’E of ISL binds (Salie et al., 2016). In addition, we speculate that S68N perturbs the binding pocket of ISL through steric hinderance; when coupled with M184V/I this may disrupt ISL binding in the NRTI pocket, which could account for the 12-fold resistance observed. The same resistance is not seen with S68G/M184V, which could be due to the smaller glycine residue present instead of a bulkier asparagine.

The hypersensitivity to ISL of RT_K65R across all subtypes examined suggests that the previously identified mechanism for hypersensitivity, a decrease in the excision efficiency of ISL-MP from the 3’ terminated primer, likely holds true in all isolates (Michailidis et al., 2013). With M184V, we see variation in resistance to ISL between isolates; however, with only one representative isolate for each subtype, we cannot associate these differences with specific subtypes. Nonetheless, all isolates containing M184V displayed some resistance to ISL and were susceptible to TDF. Similar observations were made with RT_K65R/M184V mutants: all exhibited resistance to both ISL and TDF, albeit to varying degrees. Typically, the resistance of K65R/M184V mutants was slightly reduced relative to that of the respective individual resistance mutant to its corresponding antiviral. This is presumably a consequence of the mutually incompatible resistance profiles of K65R and M184V. Interestingly, this pattern was not seen with CRF_AE isolate; rather, there was greater resistance to ISL and TDF with CRF_AE_K65R/M184V_ (9-fold resistance to ISL) compared to CRF_AE_M184V_ (3.5-fold resistance to ISL). Upon examination of the RT genome of the isolates studied, G68 was present rather than the more common S68; this mutation (S68G) was also selected during our passaging experiments. In the context of CRF_AE_K65R/M184V_, G68 appeared to confer greater resistance to ISL, relative to CRF_AE_K65R/M184V_ carrying S68. Nevertheless, the resistance observed to ISL by CRF_AE_S68G/K65R/M184V_ remained under 10-fold relative to the WT. It is also important to note that we did not see the same resistance pattern in NL4-3, suggesting that other residues (outside positions 65, 68, and 184) play a role in the resistance observed. We also see specific infectivity defect of G68 present in CRF_AE WT as compared to S68; however, in the context of K65R, M184V, K65R/M184V we saw no significant differences between Ser68 and Gly68. A previous study also found S68G in a CRF_AE isolate, where it was identified as a potential compensatory mutation for K65R (Li et al., 2020). Similarly, we found that S68G/N mutations were selected in the presence of K65R in our passaging experiments (Li et al., 2020). Our data suggest that the effects of S68G/N mutations on resistance and fitness are minor and dependent on genetic context; however, small effects may become significant during prolonged viral replication, such as may occur where ISL/TFV combinations are widely employed in HAART.

Our results highlight the high genetic barrier to resistance of TDF/ISL combinations, suggesting high efficacy for this combination if used therapeutically on a global scale. We demonstrate that TDF/ISL have opposing resistance profiles and that mutations that have been shown to confer resistance to these antivirals individually, remain susceptible to ISL and TFV in combination. Taken together these two drugs hold promise for a global combination regimen due to their high barrier to resistance, opposing resistance profiles, and efficacy against a broad range of subtypes.

## Acknowledgements

Michael Parniak was involved in the early research and writing of this study; as he is now deceased, it has not been possible to obtain his approval of the final version of the manuscript. This study was supported in part by NIH grants R37 AI076119 to S.G.S. and T32 GM008367 and F31 AI155158 (training funds for M.E.C.) and F31 AI172618 (training funds for A.S.). S.G.S. acknowledges funding from the Nahmias-Schinazi Distinguished Chair in Research. This study was supported in part by the Emory Integrated Genomics Core (EIGC), which is subsidized by the Emory University School of Medicine and is one of the Emory Integrated Core Facilities. Additional support was provided by the National Center for Advancing Translational Sciences of the National Institutes of Health under award UL1TR000454. The content is solely the responsibility of the authors and does not necessarily reflect the official views of the National Institutes of Health.

The following reagents were obtained through the AIDS Research and Reference Reagent Program, Division of AIDS, NIAID, NIH: MT-2 cells from Douglas Richman, P4-R5 MAGI cells from Nathaniel Landau. The TZM-GFPs were obtained from Massimo Pizzato (Trento University). Tenofovir disoproxil fumarate was obtained through the NIH AIDS Reagent Program. The panel of 60 international HIV-1 isolates were obtained through the AIDS Reagent Program, Division of AIDS, NIAID, NIH from the Joint UN Programme on HIV/AIDS, Dr. Victoria Polonis, Dr. Robert Gallo, Dr. Nelson Michael, and Dr. Smita Kulkarni (Brown et al. J. Virology 79(10):6089-6101, 2005). We thank Drs. Ujjwal Neogi and Anders Sönnerberg for providing HIV isolates for representative clones used in this study.

## Notes

### Competing Interest Statement

Hiroaki Mitsuya is a co-inventor of EF-dA/islatravir

